# Identification of a chromosomally-encoded sucrose operon-like gene cluster in *Vibrio parahaemolyticus* strain PH05 isolated from Negros Island, Philippines

**DOI:** 10.1101/2021.02.27.433172

**Authors:** Czarina Anne E. De Mesa, Remilyn M. Mendoza, Edgar C. Amar, Leobert D. de la Peña, Cynthia P. Saloma

**Author notes:** Contact information: Czarina Anne E. De Mesa, Remilyn M. Mendoza, Edgar C. Amar, Leobert D. de la Peña. These authors contributed equally to this work. Author for correspondence: Cynthia P. Saloma Ph.D / Loc 4703-04.

## Abstract

The ability of bacteria to metabolize a wide variety of carbon sources has been known to aid in their ability for efficient colonization. *Vibrio parahaemolyticus,* a known aquatic pathogen has been reported to have the ability to metabolize a number of carbohydrates including D-glucose, D-galactose, L-arabinose, D-mannose, and D-ribose to name a few. Classical isolation of *V. parahaemolyticus* from other members of the family *Vibrionaceae* relies on its carbon utilization pattern. Conventionally, *V. parahaemolyticus* lacks the ability to utilize sucrose and this has been the basis for its isolation using the Thiosulfate-citrate-bile salts-sucrose (TCBS) agar. Reports of *V. parahaemolyticus* having the ability to utilize sucrose have been presented yet there is paucity of information and detailed study on this phenotype. In this study, we report the *V. parahaemolyticus* strain PH05 that has the ability to metabolize sucrose. Phenotypic and genotypic characterization of this *V. parahaemolyticus* strain isolated from Negros Island, Philippines, revealed that *V. parahaemolyticus* strain PH05 is atypical appearing yellow on TCBS agar plates. It is capable of utilizing sucrose, unlike the majority of *V. parahaemolyticus* isolates. Genome analyses of this strain revealed the presence of a chromosomally encoded sucrose operon-like gene cluster encoded in chromosome 2 with the following sucrose-utilization associated genes: scrY, ccpA, treP, scrK, and scrB genes coding for sucrose porin, catabolite control protein A, PTS System sucrose-specific EIIBC component, fructokinase, and sucrose-6-phosphate hydrolase. The mode of transmission of these genes to *V. parahaemolyticus* strain PH05 is still unknown. However, the presence of insertion sequences (IS) and phage elements in the same chromosome suggests horizontal gene transfer events. Taken together, our results point to the possibility that acquired sucrose utilization genes may contribute to the fitness of *V. parahaemolyticus* strain PH05 in the environment.

## INTRODUCTION

*Vibrio parahaemolyticus* is a gram-negative halophilic bacterium usually associated with sediments, shellfish, and zooplankton in marine and estuarine environments [5, 52]. Tran and colleagues [59] identified a toxic strain of *V. parahaemolyticus* as the leading cause of acute hepatopancreatic necrosis disease (AHPND) in shrimp. Moreover, clinical strains of *V. parahaemolyticus* were identified to cause gastroenteritis accompanied by abdominal cramps and watery diarrhea when raw or undercooked seafood contaminated with *V. parahaemolyticus* is consumed [5,16].

*Vibrio* species in general metabolize D-glucose anaerobically via the Embden-Meyerhof pathway for energy production [36]. It has been established that the ability of bacteria to utilize a wide variety of unique carbohydrates and use them efficiently, aid in the effective colonization of bacteria in new niches [39]. In Freter’s nutrient-niche hypothesis, it is proposed that a bacterial species must be able to utilize a limiting nutrient better than another bacterial species for successful colonization [15]. *Vibrio parahaemolyticus* which is known to have adapted and evolved to colonize and establish niches in the intestines of humans and marine organisms such as fish and shrimps, has been reported to utilize 71 different carbon sources. Among these carbon sources are N-acetyl-D-glucosamine, mucus sugars such as L-arabinose, D-galactose, D-gluconate, D-glucuronate, D-mannose, D-glucosamine, and D-ribose [21, 39, 60]. Although sucrose-utilizing *V. parahaemolyticus* strains have been reported [26, 42], no genotypic data on this phenotype have been available.

The ability to metabolize sucrose is a variable trait in the family *Vibrionaceae,* and this has been the basis of isolation using the Thiosulfate-citrate-bile salts-sucrose (TCBS) agar [1]. TCBS is a selective and differential culture medium used in the selective isolation and cultivation of *Vibrio* species [36, 56]. Essential components of TCBS medium include yeast extract and biological peptone, that serve as the source for nitrogen and amino acids; sodium thiosulfate and sodium citrate, selective agents that render the medium alkaline, inhibiting the growth of coliforms; Ox bile, which inhibits the growth of gram-positive bacteria; sucrose, carbohydrate source; sodium chloride for optimum growth of *Vibrio* spp; ferric citrate, detects production of hydrogen sulfide from sodium thiosulfate utilization; bromothymol blue, a pH indicator; and agar as a solidifying agent. Conventionally, colonies of *Vibrio parahaemolyticus* strains in TCBS agar appear blue to green and do not ferment sucrose [56]; hence no acid production would entail changing TCBS agar from green to yellow. The US Food and Drug Administration uses this biochemical reaction to identify strains of *V. parahaemolyticus* [23].

Sucrose, composed of one glucose unit and one fructose unit, is the most abundant disaccharide in the environment due to its origin in higher plant tissues [40]. Metabolism of sucrose involves the catalytic activities of sucrose-6-phosphate hydrolases and sucrose phosphorylases.

This study reports an atypical *V. parahaemolyticus* strain PH05 isolated in Negros, Philippines, under the Philippine Shrimp Pathogenomics Program (PSPP) of the Philippine Genome Center. Bioinformatics analyses of its draft whole-genome sequence revealed the presence of a sucrose operon-like gene cluster that has not been reported previously in *V. parahaemolyticus* strains.

## MATERIALS AND METHODS

### Isolation of *V. parahaemolyticus* strains

*Vibrio* isolates from the Philippine Shrimp Pathogenomics Program (PSPP) collection of the Philippine Genome Center were isolated in 2014-2016. The hepatopancreas of five (5) shrimp were dissected and placed in a tear-resistant sterile plastic bag for homogenization using normal saline solution (NSS). Serial dilutions from 10^-1^ to 10^−4^ were spread plated in Nutrient Agar *(Pronadisa)* with 1.5% NaCl (NAN), Thiosulfate Citrate Bile Salts Sucrose (TCBS) agar *(Pronadisa),* and *Vibrio* chromogenic agar *(Pronadisa)* in duplicates. The presence of luminous bacteria was noted, and dominant colonies were isolated after 18-24h. Pure cultures were ascertained before preparing 20% glycerol stock and storage at −80 °C biofreezer for biobanking.

*Vibrio* spp. from PSPP isolates initially identified as *V. parahaemolyticus* were randomly selected for revival. From glycerol stock, a 100 μl aliquot was used for inoculation in 5 ml nutrient broth with 1.5% sodium chloride (NBN), and tubes were then incubated overnight at 37 °C. Turbid broths were then streaked on TCBS agar plates and incubated at 37 °C for the selective growth of *Vibrio* species. Single colonies from these TCBS plates were streaked on NAN for further Gram-staining procedure, colony morphology observation, and biochemical characterization.

### Genome Assembly and Annotation

The Philippine Shrimp Pathogenomics Program of the Philippine Genome Center previously sequenced more than 400 bacterial pathogens of shrimp as previously described [35]. All analyses were done using default settings unless otherwise stated. The whole genome sequence of *V. parahaemolyticus* strain PH05 was initially pre-processed using *Fastp v0.20.0* [8] and assembled using *SPAdes genome assembler v3.11.1* [32]. Assembled contigs were assessed using *QUAST v5.0.2* [30] and filtered using *SeqKit* [50] such that contigs with less than 1000 bases were omitted. Assembled and filtered contigs were then annotated using *Prokka v1.13.7* [48], Rapid Annotation using Subsystem Technology (RAST) [4, 6, 34], and *eggNOgmapper* [18]. The presence of plasmids was queried through *PlasmidSpades* [32], *plasmidFinder* [9], and *PLSDB* database [16]. A reference-based assembly of contigs was done using *chromosomer* with a ratio threshold of 1 [55]. Identification of IS elements, transposons, and phage elements was made by comparing the draft genome against the sequences found in *ISfinder* [51], *ICEfinder* [28], and *PHASTER* [3].

### Confirmation of the identity of *V. parahaemolyticus* strain PH05

Taxonomic affiliation as *Vibrio parahaemolyticus* was confirmed by Average Nucleotide Identity (ANI) calculation implemented in *pyani* v0.2.9 [38] using default settings against the representative genomes of *Vibrio parahaemolyticus, Vibrio harveyi, Vibrio campbellii* and *Vibrio alginolyticus.* The *16s rRNA* gene of *V. parahaemolyticus* strain PH05 was also extracted from the whole genome sequence and was subjected to BLASTn (wwww.blast.ncbi.nlm.nih.gov) analysis.

### Gene Finding

Genes related to sucrose utilization such as: sucrose porin, catabolite control proteins, PTS System sucrose/threhalose-specific, fructokinase, sucrose-6-phosphate hydrolase, and invertase [40] were searched from the annotated genomes of *V. parahaemolyticus* strain PH05, as well in *V. parahaemolyticus* PH 1339 [35] and *V. parahaemolyticus* RIMD2210633 [59] which served as the controls. The genomes of *V. parahaemolyticus* strain PH05 and *V. parahaemolyticus* strain PH1339 were compared using Mauve [12]

## RESULTS AND DISCUSSION

### *V. parahaemolyticus* strain PH05 showed an atypical phenotype in TCBS agar

*V. parahaemolyticus* is associated with food-borne diseases after seafood consumption and recently, with acute hepatopancreatic necrosis disease (AHPND), a devastating infection of cultured shrimp [27], Isolation of *V. parahaemolyticus* commonly relies on the use of TCBS, a medium selective against gram-positive bacteria and differentiates sucrose-utilizing *V. cholerae* from non-sucrose utilizing *Vibrio* like *V. parahaemolyticus* [33]. However, the colonies of the putative *V. parahaemolyticus* strain PH05 on TCBS agar were round with an entire margin and notably colored yellow, indicating acid production from sucrose utilization (Figure 1). This characteristic, while highly atypical in *V. parahaemolyticus,* had been previously reported [26, 42].

**Figure 1.**
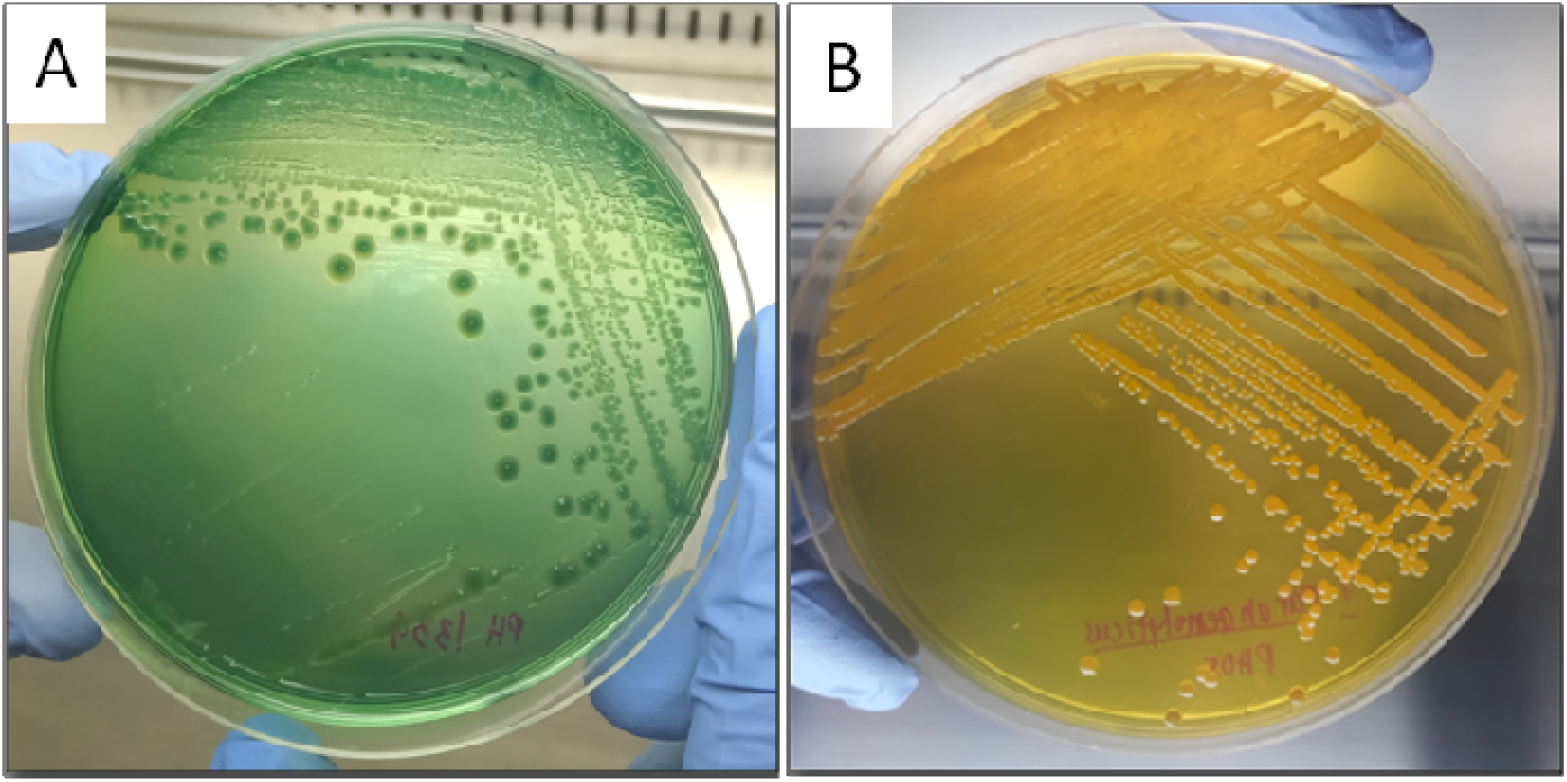
Plate cultures (24-h) of V. parahaemolyticus strain PH1339 and the atypical V. parahaemolyticus strain PH05 in TCBS agar. (A) *Vibrio parahaemolyticus* strain PH1339 shows the typical green to bluish-green colonies in TCBS agar. (B) Yellow colonies of putative *Vibrio parahaemolyticus* strain PH05 in TCBS agar indicative of sucrose utilization.

The biochemical characteristics of putative *V. parahaemolyticus* strain PH05 initially obtained from API 20NE analysis are presented in Table 1. Inputting this given biochemical profile and the ability of the putative *V. parahaemolyticus* strain PH05 to utilize sucrose in the ABIS online laboratory tool for bacterial identification database [11] resulted in the identification of the putative *V. parahaemolyticus* to *Aeromonas media* (97.2%). However, since this biochemical identification is not enough and not conclusive, we proceeded to the verification of the identity of the putative *V. parahaemolyticus* strain PH05 through molecular methods.

**Table 1.** Biochemical characteristics of putative *V. parahaemolyticus* strain PH05

| **Reactions/Enzymes** |  |
| --- | --- |
| Reduction of Nitrates to nitrite and nitrite to nitrogen | + |
| Indole production | + |
| Glucose fermentation | + |
| Arginine dihydrolase | - |
| Urease | - |
| Hydrolysis (B-glucosidase) Esculin | + |
| Hydrolysis (protease) gelatin | + |
| B-galactosidase | + |
| **Assimilation:** |  |
| Glucose | + |
| Arabinose | + |
| Mannose | + |
| Mannitol | + |
| N-acetyl glucosamine | - |
| D-maltose | + |
| Potassium gluconate | + |
| Capric acid | - |
| Adipic acid | - |
| Malic acid | + |
| Trisodium citrate | - |
| Phenylacetic acid | - |
| Oxidase | + |

### Identification of of *Vibrio parahaemolyticus* strain PH05

Raw reads of the whole genome sequence of the putative *V. parahaemolyticus* strain PH05 were assembled and annotated. The draft genome assembly of the putative *Vibrio parahaemolyticus* strain PH05 resulted in 26 contigs with a total length of 4954268 bp and G + C content of 45.37%. The N50 of the assembly was 729047 bp. Annotation with *Prokka v1.13.7* [48] revealed 4442 coding sequences (CDS), 110 tRNA, 43 miscellaneous RNA, and one tmRNA.

The python module *pyani* v0.2.9 [32] was used to calculate the ANI using the genome sequence of *V. parahaemolyticus* strain PH05 against the representative genomes of *Vibrio parahaemolyticus, Vibrio harveyi, Vibrio campbellii,* and *Vibrio alginolyticus.* Blast [8] and MUMmer [29] were both used in the calculation of ANI similarity percentages. The ANIb (98.43%) and ANIm (98.55%) percentage identities against the genome sequence of *V. parahaemolyticus* RIMD2210633 confirmed the identity of *V. parahaemolyticus* strain PH05 (see Table 2 & Figure 2, supplementary material). Although there is a difference in values of ANIb and ANIm, the 95-96% values of each method correspond to the 60-70% similarity cut-off point for bacterial species delineation of DNA-DNA hybridization method [2, 24]. The result of ANI calculations supports its initial identification using Blastn analysis of the partial sequence of *16s rRNA* gene (See Table 1 Supplementary Material). However, with only ~60% of the *16s rRNA* sequence recovered, its utility for accurate strain identification is limited [20]. Furthermore, highly similar *Vibrio* species (such as *V. cholerae* and *V. mimicus)* cannot be differentiated with their 16s rRNA gene sequences alone. [31]. The importance of using ANI in bacterial identification is especially highlighted in this study because traditional protocols (TCBS culture, Biochemical tests) we used together with *16s rRNA* gene were insufficient in identifying the sucrose-fermenting *V. parahaemolyticus* strain PH05.

**Figure 2.**
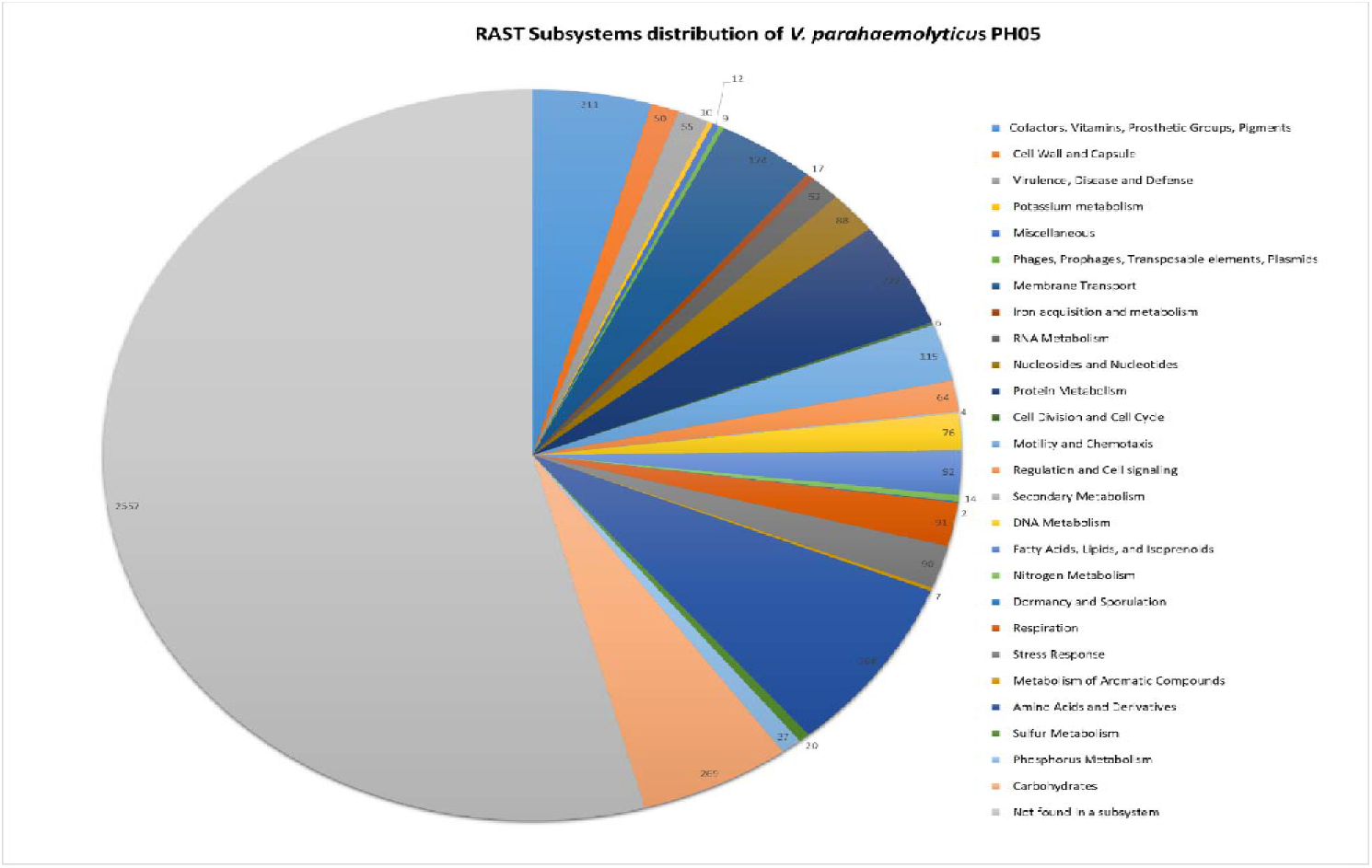
Distribution of subsystems in *V. parahaemolyticus* strain PH05 identified by RAST. Genes involved in sucrose utilization were identified and classified under the carbohydrates subsystems

**Table 2.**
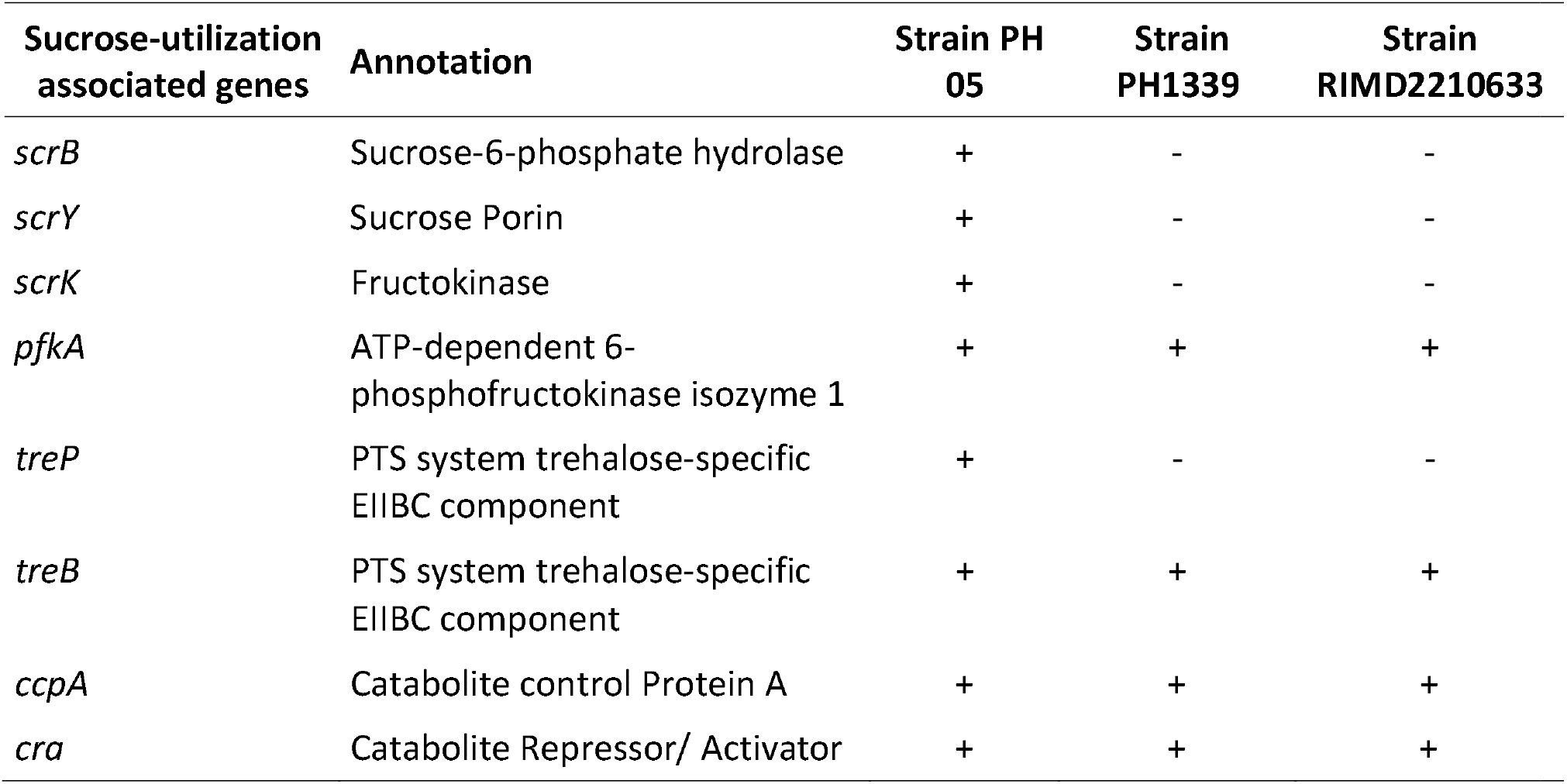
Sucrose-utilization-associated genes found in the annotated genome of *Vibrio parahaemolyticus* strain PH05.

### Presence of a sucrose operon-like gene cluster in chromosome 2 of *Vibrio parahaemolyticus* strain PH05

Annotation by RAST identified 4716 genes in V. *parahaemolyticus* PH05, to which 2557 (54%) were not categorized to any subsystem and 2159 (46%) genes were classified under different RAST subsystems (Figure 2). Subsystems are defined as functional roles that together implement a specific biological process or structural complex [4, 6, 34]. Among the notable subsystems, 55 genes were identified related to Virulence, Disease, and Defense; Phages, Prophages, Transposable elements, and Plasmids subsystem; while in the carbohydrates subsystem, 269 genes were identified including sucrose utilization genes.

To determine if the sucrose utilization gene is unique in *V. parahaemolyticus* strain PH05, The annotated genomes of *V. parahaemolyticus* strain PH05, *V. parahaemolyticus* strain PH1339, and *V. parahaemolyticus* RIMD2210633 were both searched for genes related to sucrose metabolism such as sucrose-6-phosphate hydrolase, sucrose porin, PTS System-sucrose/trehalose-specific, fructokinase, sucrose permease, and sucrose phosphorylase [39]. It is noteworthy that the genomes of the *V. parahaemolyticus* strains scanned all possessed genes encoding ATP-dependent fructokinase, catabolite control protein, and trehalose-specific PTS system. In contrast, only *V. parahaemolyticus* strain PH05 genome contained genes coding for a sucrose porin and sucrose-6-phosphate hydrolase (Table 2). Visualization in *Artemis* [43] revealed that *V. parahaemolyticus* strain PH05 has a sucrose operon-like gene cluster (Figure 3). This gene cluster indicated the ability to utilize sucrose as a carbon source through PTS-dependent sucrose transport and catabolism.

**Figure 3.**
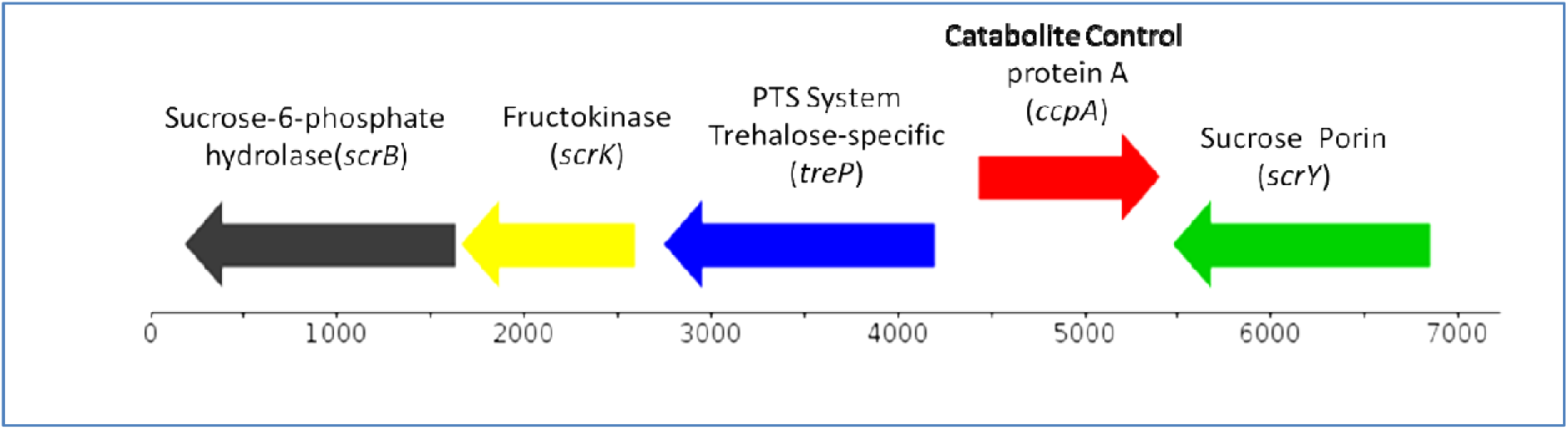
Sucrose operon-like gene cluster found in V. parahaemolyticus strain PH05. *V. parahaemolyticus* strain PH05 is atypical as it possesses a putative sucrose operon that consists of genes coding for sucrose-6-phosphate hydrolase, fructokinase, and PTS System which are the key players in a PTS-dependent sucrose transport and metabolism. A sucrose specific porin was also evidently found localized with the putative sucrose operon, as well as the catabolite control protein A that belongs to the LacI-GalR family of transcriptional regulators that has been reported to be often associated with sucrose-utilization genes. All the genes in the cluster are in the reverse stand except for the ccpA which is found on the forward strand.

In PTS-dependent sucrose transport and catabolism, external sucrose is transported and phosphorylated by the PTS system’s three-enzyme complex, giving off phosphorylated sucrose, sucrose-6-phosphate, to the cell. This is hydrolyzed by sucrose-6-phosphate hydrolase, giving off glucose-6-phosphate that can directly enter the glycolytic pathway; and fructose that is phosphorylated by fructokinase to fructose-6-phosphate that also enters the glycolytic pathway [40]

The amino acid sequence of the sucrose-6-phosphate hydrolase (s*crB*) of *V. parahaemolyticus* strain PH05 contains the signature pentapeptide β-fructosidase motif (NDPNG) (Supplementary material, Figure 2), which is found to be part of a large 14-amino acid motif known as the sucrose-binding box wherein the aspartate residue act as the active site [40,19]. Meanwhile, sucrose porin (s*crY*) is an example of a specific porin that act as channels across the membrane [53]. Passage of sucrose through the sucrose porin has been hypothesized to initially begin with sucrose molecules diffusing from the external solution to a “trapping zone” in the sucrose porin. Then sucrose molecules slide along a greasy slide; enter and pass the binding region to enter the cell [14, 53].

Blastx analysis of the *treP* and *ccpA* genes revealed that the *treP* gene has 99.79% identity to the PTS system sucrose-specific IIB component (Glc family) /PTS system sucrose-specific IIC component (Glc family) of *V. parahaemolyticus* in the database; while the *ccpA* gene codes for a LacI family transcriptional regulator, catabolite control protein A, that is 98.48% identical to that of *V. parahaemolyticus* in NCBI database. LacI-GalR family of transcriptional regulators has been mentioned to be often associated with sucrose-utilization genes [40], inducing or repressing the structural genes in a substrate-specific manner for better allocation or utilization of energy by the cell.

It was initially hypothesized that the sucrose gene cluster found in *Vibrio parahaemolyticus* strain PH05 was extra-chromosomally encoded since there were reports of conjugative plasmid pUR400 found in *Salmonella* and *E. coli* [61, 44, 45] capable of transfer of sucrose utilization genes. To confirm this, putative plasmid sequences were assembled using *PlasmidSpades* [32], and plasmid-related elements were searched in *plasmidFinder* [9] and the *PLSDB* database [16]. However, no plasmids were detected in the draft genome. It was therefore highly probable that the sucrose genes were chromosomally encoded. The draft genome was then scaffolded into two chromosomes using the tool *chromosomer* [55], which uses genome similarities with a reference genome. The reference genome *V. parahaemolyticus* RIMD2210633 was first used as a reference, but only the first chromosome was assembled (~3 Mb). Unexpectedly, we found a complete genome of *V. parahaemolyticus* strain HA2, which possessed chromosomally encoded scr genes since it appeared in the result of the blastn and blastx analyses of the sucrose porin and sucrose-6-phosphate hydrolase. The result of assembly using *V. parahaemolyticus* strain HA2 as reference resulted in two chromosomes (Figure 4) where the scr genes were found to be encoded in the smaller chromosome 2 (~1.7 Mb). The assembly of the draft genome into the two chromosomes points to the high similarity between the genome of *V. parahaemolyticus* strain HA2 and *V. parahaemolyticus* strain PH05, which are both aquaculture isolates.

**Figure 4.**
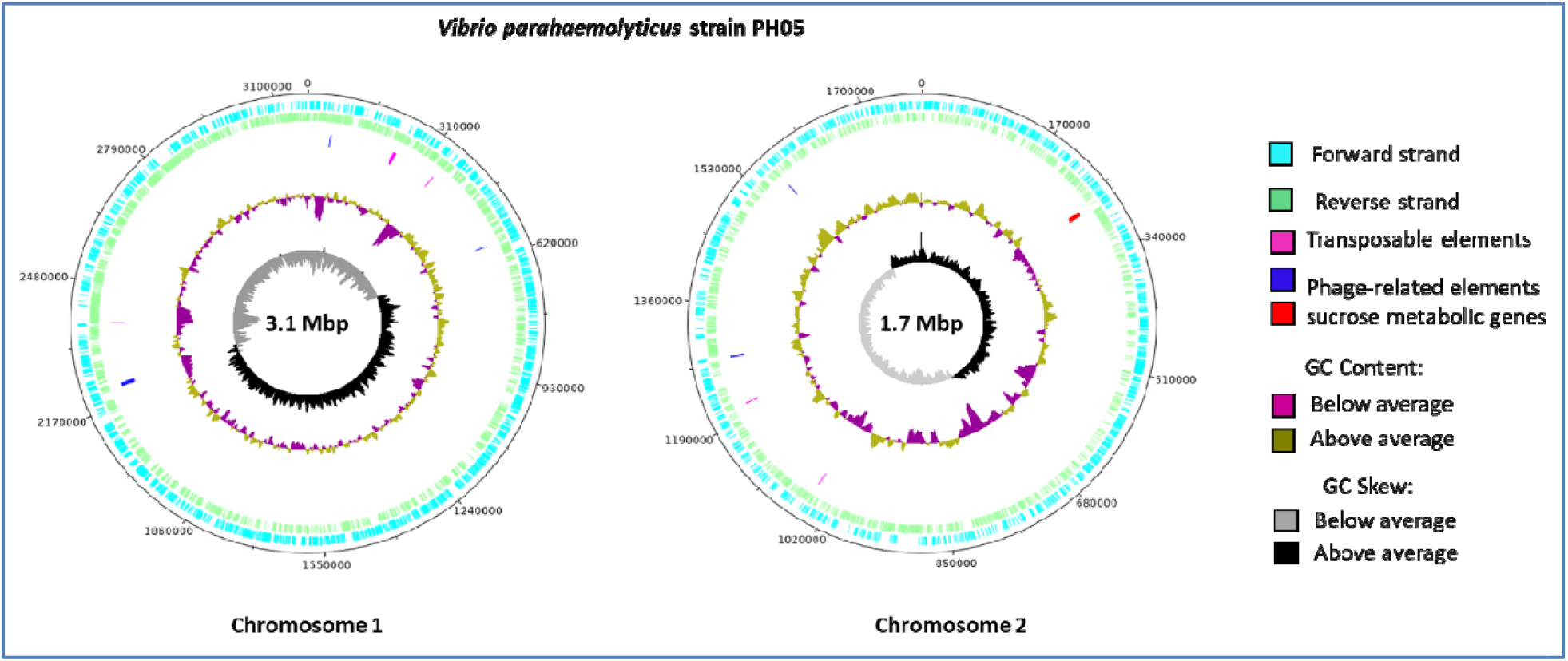
Genome organization of V. parahaemolyticus strain PH05. The two (2) chromosomes of *V. parahaemolyticus* strain PH05 (mapped against *V. parahaemolyticus* HA2) showing the genome composition and the location of the sucrose operon like gene cluster in the chromosome 2, as well as relevant elements for HGT. The figure was generated using DNA plotter of the visualization tool Artemis [43]

We then compared, the segment of the chromosome of *V. parahaemolyticus* strain PH05 containing the sucrose-operon-like cluster to that of *V. parahaemolyticus* HA2 and *V. parahaemolyticus* strain PH1339. Based on genome comparison (Figure 5), it appeared likely that there is an insertion of sucrose metabolic genes in the chromosome of *V. parahaemolyticus* strain PH05 as well as in *V. parahaemolyticus* strain HA2 that is not found in *V. parahaemolyticus* strain PH1339. This is supported by the presence of genes flanking the sucrose operon-like gene cluster in the chromosomes of both *V. parahaemolyticus* strains PH05 and HA2. The apparent shift of the genes and absence of the *yyaP* gene in the chromosome segment containing the sucrose operon-like gene cluster in *V. parahaemolyticus* strain PH05 was also evident. *V. parahaemolyticus* strain PH1339, similar to strain PH05, was isolated in *Penaeus vannamei.*

**Figure 5.**
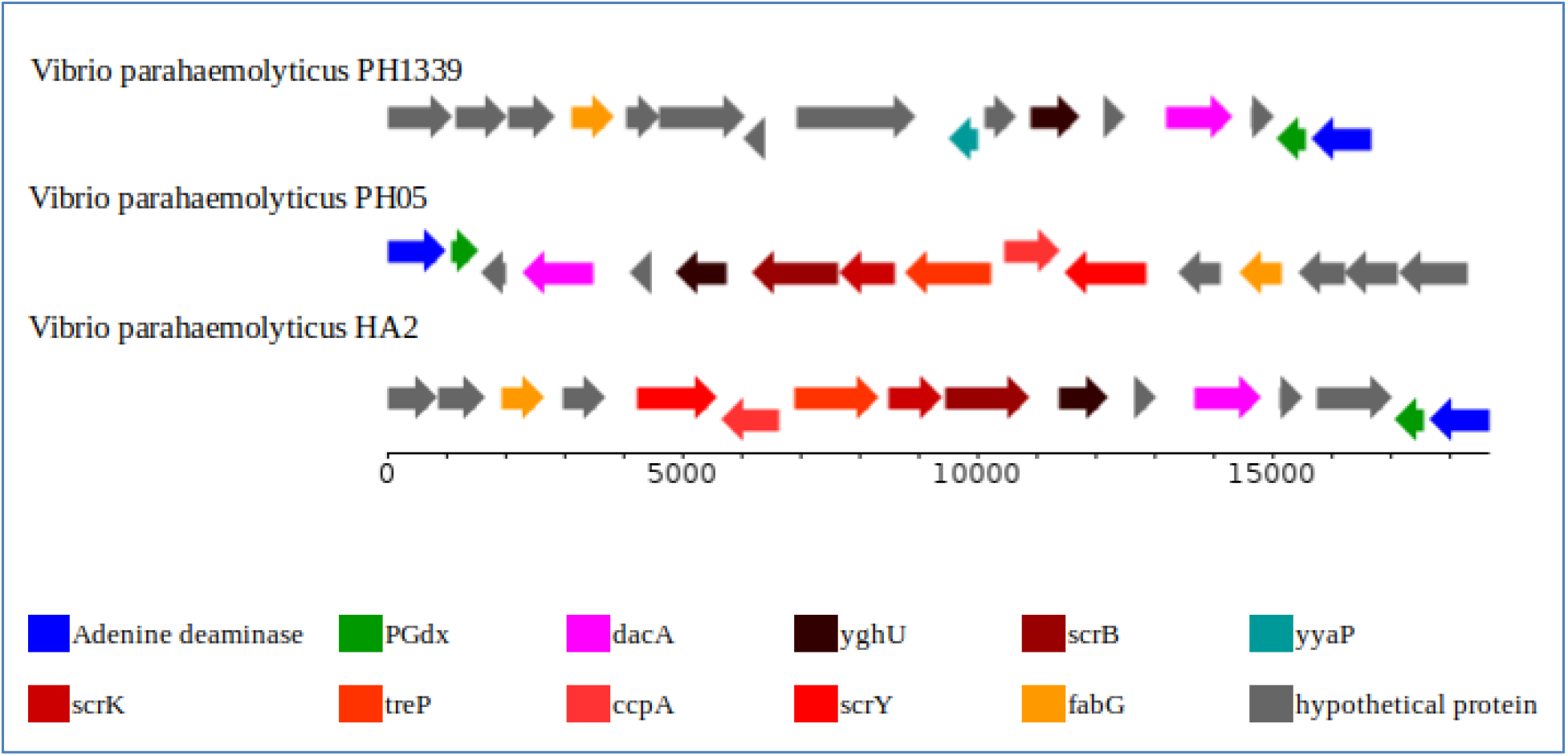
Linear representation of the Chromosome 2 segment of V. parahaemolyticus strain PH05 harboring the sucrose operon-like gene cluster. Highly similar gene architecture of *V. parahaemolyticus* strain PH05 was mapped with that of *V. parahaemolyticus* strain HA2 (Accession: SAMN07680340). Evident insertion of the sucrose metabolic gene cluster in *V. parahaemolyticus* strain PH05 was also observed when compared with *V. parahaemolyticus* strain PH1339 (Accession: SRR9937595). Genome comparison (supplementary material Figure 3 and 4) was visualized using Mauve [12].

To further investigate the mechanism of insertion of sucrose-utilization genes in chromosome 2 of *V. parahaemolyticus* strain PH05, we looked at several databases and annotation tools. *Prokka* [48] was able to annotate a few transposons that were all found in chromosome 1. Amino acids from the genome were functionally annotated using eggNog mapper [18] and revealed some hypothetical proteins in chromosome 2 as transposases. ISfinder [51] was used to identify insertion sequences usually associated with transposons. Phage elements were also identified through *Prokka* [48] and *PHASTER* [3]. However, these were found to be localized far from the scr gene cluster. Taken together, the scr gene cluster was probably transferred through a natural transformation event since *Vibrio* species are known to be naturally competent [54]. For instance, Chitin-induced competence in *V. cholerae* allowed it to acquire gene clusters that altered its serogroup [7].

On the other hand, foreign DNA can be integrated into the chromosome by illegitimate recombination in which even short homologous regions facilitate the integration of a foreign DNA [14]. In the mauve comparisons, there were homology sites with the *V. parahaemolyticus* strain HA2 with the sequences encoding the scr gene cluster of *V. parahaemolyticus* strain PH05 (Figure 3 in the supplementary). These sites were probably used to facilitate the integration of the scr gene clusters during its period of competence.

The ability to metabolize a wide array of carbon sources confers an adaptive advantage to the organism in an environment where glucose is limiting. Sucrose, a disaccharide consisting of one glucose and one fructose unit, is the most abundant disaccharide owing to its origin in higher plant tissues [40, 49], and not all microorganisms possess the machinery to metabolize this carbohydrate.

One way by which *V. parahaemolyticus* is differentiated from other *Vibrio* species such as *V. cholerae* and *V. alginolyticus* is through their difference in colony morphology in TCBS agar plates. It has been established that *V. cholerae* and *V. alginolyticus* are sucrose fermenters, hence, the yellow colonies that they produce in TCBS agar plates, while *V. parahaemolyticus* is known to be unable to ferment sucrose and exhibits green colonies in TCBS agar. However, previous studies have reported yellow *V. parahaemolyticus* isolates from oysters and mussels [42] and from the diarrheal stool of a patient [26]. In the present study, we report an atypical *V. parahaemolyticus* strain PH05 where a sucrose operon-like gene cluster was found encoded in its chromosome 2.

*Vibrio parahaemolyticus* strain PH05 was isolated from *Penaeus vannamei* from a shrimp pond in Negros Island, Philippines. Its ability to utilize sucrose may be a form of metabolic adaptation to thrive in that specific environment. In some aquaculture ponds, molasses (which is high in sucrose) was used as an organic fertilizer [47] and as a growth promoter of heterotrophic bacteria to control the accumulation of nitrogenous wastes [46]. In the study of Tendencia, Bosma, & Sorio [57], molasses was used to promote the growth of “yellow *Vibrios”* to act against pathogenic *Vibrio,* rendering shrimp less susceptible to Whitespot Syndrome Virus (WSSV). Antagonistic action of these yellow *Vibrio* against pathogenic green *Vibrio* species such as *V. parahaemolyticus* and *V. harveyi* can be attributed to bacteriocin production [58]. However, the ability to metabolize sucrose in an environment with adequate sucrose concentrations may have also contributed to their ability to compete with non-sucrose fermenting *Vibrio* species. The ability of *V. parahaemolyticus* strain PH05 to utilize sucrose may add to its overall fitness given that in its environment, sucrose is a limiting nutrient [39].

In *Streptococcus pneumoniae,* a gram-positive bacterial pathogen, two sucrose-metabolizing systems, the *sus* and scr systems, were reported and described by Iyer and Camilli [19] to play niche-specific roles in virulence. Sucrose is a major carbon source used by S. *pneumoniae* during colonization and infection. Their study found that the uptake and catabolism of sucrose have essential roles in colonization of murine nasopharynx to which the *scr* system is primarily involved, and an accessory role in the infection of the lungs with the *sus* system as a key player. Moreover, in another study by Shelburne and colleagues [49], the catabolite control protein A *(ccpA)* ortholog in Group A *Streptococcus* (GAS) was found to influence the transcript levels of many carbohydrate utilization genes and several GAS virulence factors, including the cytolysin streptolysin S, and speB (cysteine protease). Their data demonstrated that the ccpA of GAS regulates virulence factors’ production depending on the nutrient availability in the environment. In a nutrient-rich environment, the production of streptolysin S is repressed, while in a nutrient-limiting environment, the production of SpeB is increased. The interconnection between carbohydrate metabolism and pathogenesis in the gram-negative *V. parahaemolyticus* stain PH05 is an exciting aspect that can be explored and studied in the future.

## CONCLUSION

*Vibrio parahaemolyticus* strain PH05 isolated from Negros Island, Philippines in the summer of 2014, exhibits an atypical phenotype as it possesses the ability to utilize sucrose as demonstrated by its growth in TCBS. This phenotype was supported by the presence of genes associated with sucrose metabolism such as sucrose porin, sucrose-6-phosphate, fructokinase, PTS system sucrose-specific IIC component (Glc family), and catabolite control protein A. These genes are arranged as a sucrose operon-like gene cluster found in chromosome 2 (~1.7 Mb). The presence of IS sequences, transposases, and phage elements suggests horizontal gene transfer events in the chromosome with the scr gene cluster’s acquisition a likely metabolic adaptation to the environment possibly linked to its virulence and pathogenicity.

## CREDIT AUTHOR STATEMENT

**De Mesa, CA** and **Mendoza, RM** were involved in Conceptualization, Methodology, Investigation, Writing (original draft), and Visualization; **Amar, EA** and **De la Peña, LD** were involved in the Conceptualization, Collection of Specimen, Biochemical Characterization and Funding acquisition; **Saloma, CP** was involved in Conceptualization, Writing (Review and Editing), Resources, Supervision and Funding acquisition.

## DECLARATION OF COMPETING INTEREST

The authors declare no conflict of interest.

## ACKNOWLEDGMENT

The authors would like to acknowledge the contributions of Ms. Sarah Penir and the people who previously worked under the Philippine Shrimp Pathogenomics Program, the Core Facility for Bioinformatics of Philippine Genome Center and the Negros Prawn Producers Cooperative for their generosity and assistance.

## FUNDING

This work was supported by a Department of Science and Technology, Philippine Council for Agriculture, Aquatic and Natural Resources Research and Development (DOST-PCAARRD) grant to CPS, EAA and LDD and a DOST Science Education Institute fellowship to CADM and RMM.

## NUCLEOTIDE SEQUENCE DATA

The Illumina reads of *V. parahaemolyticus* PH05 have been deposited under the accession no. SRR13286682, and the genome assembly has been deposited at DDBJ/EMBL/Genbank under the accession no. JAELVO000000000.

## Abbreviations

TCBS: Thiosulfate-citrate-bile salts-sucrose
NAN: Nutrient Agar with 1.5% NaCl
NBN: nutrient broth with 1.5% NaCl

## Supplementary Material

**Table 1.** Blastn result of partial *16s rRNA* gene of *Vibrio parahaemolyticus* strain PH05 showing 100% identity with *Vibrio parahaemolyticus* strains in the database.

| Description | Accession no. | Percent identity | E value | Max score | Total score | Query Cover |
| --- | --- | --- | --- | --- | --- | --- |
| *Vibrio parahaemolyticus* strain 2012AW-0154 chromosome 1, complete sequence | CP035701.1 | 100% | 0.0 | 1737 | 19031 | 100% |
| *Vibrio parahaemolyticus* strain FDAARGOS_662 chromosome 2, complete sequence | CP044071.1 | 100% | 0.0 | 1737 | 19018 | 100% |
| *Vibrio parahaemolyticus* strain MVP1 chromosome 1, complete sequence | CP043421.1 | 100% | 0.0 | 1737 | 18743 | 100% |
| *Vibrio parahaemolyticus* strain Vb0624 chromosome 1, complete sequence | CP041202.1 | 100% | 0.0 | 1737 | 19081 | 100% |
| *Vibrio parahaemolyticus* strain 160807 chromosome 2, complete sequence | CP033142.1 | 100% | 0.0 | 1737 | 1737 | 100% |

**Table 2.**
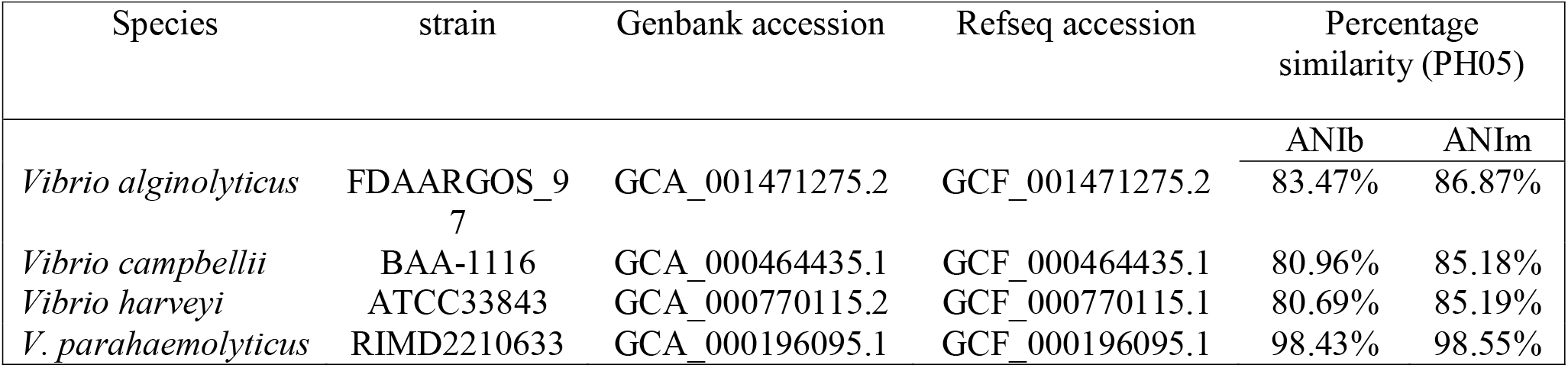
Percentage Identities of *V. parahaemolyticus* strain PH05 against *Vibrio* reference genomes.

**Figure 1.**
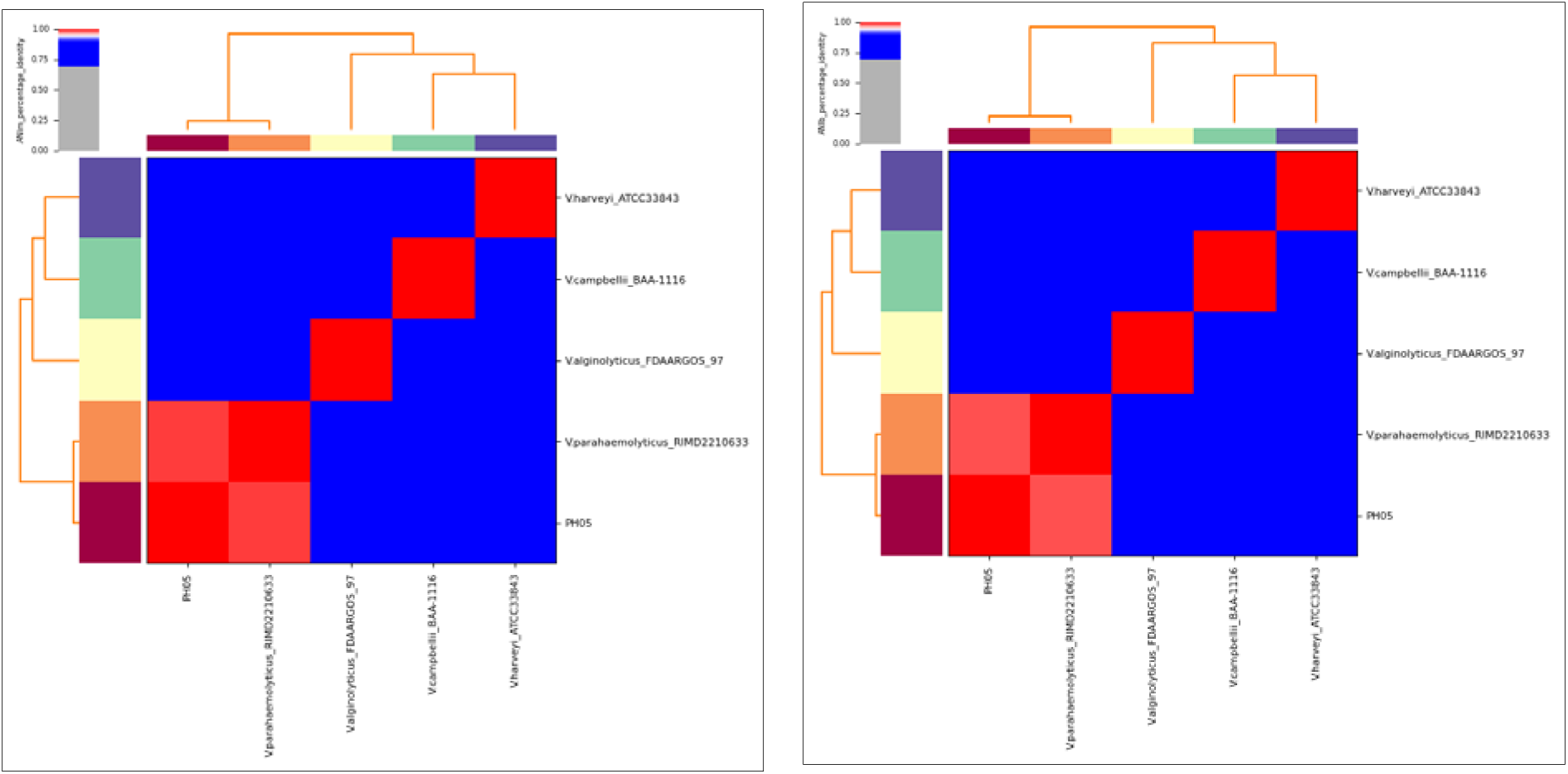
Average Nucleotide Identity (ANI) heat maps. The degree of similarity between putative *Vibrio parahaemolyticus* strain PH05 and other Vibrio species was assessed using ANI. The ANI heat maps showed higher percentage identity (98%, see Table 2) of *V. parahaemolyticus* strain PH05 with that of *V. parahaemolyticus* RIMD 2210633, confirming the identity of *V. parahaemolyticus* strain PH05. Heatmaps from both ANIb (left) and ANim (right) calculations are identical due to small differences in their percentage values.

## Notes

### Competing Interest Statement

The authors have declared no competing interest.

